# 3D Agent-based modeling without chemical signaling recreates collective behaviors seen in *Myxococcus xanthus* colonies

**DOI:** 10.1101/2025.05.23.655808

**Authors:** Katherine Copenhagen, Matthew Black, Joshua W. Shaevitz

**Affiliations:** Lewis-Sigler Institute of Integrative Genomics, Princeton University; Joseph Henry Laboratory of Physics, Princeton University

## Abstract

*Myxococcus xanthus* is a soil-dwelling bacterium that lives in dense populations of millions of cells and displays an array of collective behaviors and macroscopic patterns in response to environmental conditions. The transitions between macroscopic patterns formed by this species are driven by changes in cell motility such as increasing Péclet number via increasing cell speed and reversal period. While cells are capable of passing contact-mediated signals that can trigger reversal events, we set out to study which aspects of *M. xanthus* collective behavior may be driven by simple individual cell behavior changes in the absence of cell signaling. We do this through 3D agent-based simulations with a bead-spring chain model. We find that when tuned to match single cell properties, this model captures multicellular structures observed at both intermediate and global scales in *M. xanthus* experiments. Increasing cell speed and reversal period together (as cells do when they starve), maintains a constant overall nematic structure in the population, while significantly increasing the net flow into and out of topological defects, which drives an increased aggregation that leads to fruiting body formation during starvation.

## I. INTRODUCTION

Dense populations of reversing self-propelled rods are a class of active materials, prevalent in nature from bacterial colonies to sea algae. These collectives display a wide variety of unique behaviors and patterns due to their combination of polar and nematic features[1]. Microscopic interactions between self-propelled rods are typically nematic, i.e. head-tail symmetric, driven by steric collisions and volume exclusion as rods tend to align their long axes when packed together. Their direction of self-propulsion, however, endows particles with a natural polarity. Because of this, groups of self-propelled rods moving in the same direction create local polar order through polar sorting[2] and Fisher wave propagation[3, 4]. This accumulation of local polar order has been shown to create both traveling waves in groups of swimming bacteria[3] and, in *Myxococcus xanthus*, accumulation of local stresses leading to the out-of-plane extrusion of cells [5].

Local polar order, however, is interrupted when some fraction of the cells in a group reverse their direction of motion. At the single cell level, directional reversals offer a means of modulating their effective diffusivity [6, 7]. Tuning the period at which such reversals take place provides an internally controllable timescale by which cells can “switch” between ballistic and diffusive motion [8]. Such directional reversals appear to be exploited throughout the biological world. Neural progenitor cells have been shown to exploit reversals to promote cell aggregation [9]. Several marine species of bacteria and algae self-propel by flagellar-mediated swimming while undergoing periodic directional reversals [10, 11]. Terrestrial bacteria, which move by either pilus-based twitching or focal adhesion-based gliding, also may exhibit directional reversals[12–15].

The soil bacterium *M. xanthus* is an ideal model organism for studying the relationship between single cell properties and population level collective behaviors and pattern formation in systems of self-propelled reversing rods. Single *M. xanthus* cells are about 0.5µm wide and 5µm long and glide across surfaces at a speed of about a body length per minute, reversing their direction of motion every ∼ 8 minutes. On the other end of the scale, in populations of millions of cells, *M. xanthus* exhibits a complex multicellular lifecycle, forming different collective 3D structures depending on environmental conditions[16]. For example, groups of *M. xanthus* cells predate on other bacteria and exhibit “rippling” waves, nutrient-starved populations aggregate into macroscopic, spore-filled fruiting bodies, while well-fed populations spread out into a thin swarming layer of cells coating a surface[17]. Transitions between these different population structures are driven by changes in motility at the single cell level[18] which may be triggered by local cell-cell contacts [19] or individual cells sensing local nutrient conditions and internal state [20]. Understanding the full physical mechanisms connecting single cell behavior to population-level structure formation will increase our knowledge of *M. xanthus* life-cycle and also reveal simple single particle properties that may be exploited to drive active materials between different macroscopic phases.

Individual cells respond to nutrient stress by both moving faster and increasing the time between reversal events (reversal period). At the population level, these motility changes cause an apparent spinodal decomposition from a thinly-spread layer to a collection of well separated fruiting bodies [18]. Recent work shows that, on an intermediate length scale, well-fed cells form a thin swarm and pack together, aligning their long axes over a range of several cell lengths. As cells glide smoothly past each other within the cell layers, they exert pushing forces along the cell alignment direction. The population thus forms an active nematic material, and internal patterns form where cells become misaligned at sites called topological defects[21, 22].

During the initial stages of fruiting body formation, cells begin to climb on top of other cells, driven by the formation of new layers and holes at topological defects[22]. The stochastic accumulation of local polar order at +1/2-charge defects gives rise to a local stress accumulation that ultimately extrudes cells out of their original 2D layer and into a multi-layered structure [5]. We aim to investigate whether altering cell motility parameters without invoking cell signaling is sufficient to recapitulate the collective behaviors observed in *M. xanthus* during starvation.

Hydrodynamic modeling of active nematic materials reveals mean-field forces, such as activity, friction and pressure, which accurately predict the average flows seen in experiments around topological defects[5]. However, hydrodynamic modeling of active nematic materials may miss important features that arise from the discrete nature of the experimental systems which can be captured by agent-based simulations[23]. We use a bead-spring chain agent-based model to simulate *M. xanthus* colonies and independently tune cell speed and reversal rate. We do not add additional cell signaling or modulate any other aspects of the cell physiology which could occur in experiments that use drugs or mutants to modulate cell motility. Our model captures the collective behaviors observed experimentally in *M. xanthus* while allowing us to independently tune cell speed and reversal period.

## II. METHODS

### A. Experiment

*M. xanthus* cells were grown according to standard protocols. Wild-type cells (strain DK1622) were first taken from frozen stocks and struck onto 1.5% agar-CTTYE plates. CTTYE is a nutrient rich media, commonly used to culture *M. xanthus*, made of 1% w/v Peptone, 0.5% w/v yeast extract, 10 mM Tris-HCl (pH 8.0), 1 mM KH_2_PO4, and 8 mM MgSO_4_. Plates were incubated at 32^◦^C until the first appearance of colonies (∼ 3 − 4 days). Colonies were then inoculated into liquid media and incubated overnight, shaking at 32^◦^C. Fresh 1.5% agarose pads filled with rich media were prepared fresh for each experiment and allowed to set before a sample of cells from the overnight liquid culture was spotted onto it. After allowing the spot to dry, this pad was moved to the microscope for imaging.

Cells were imaged by confocal reflected laser light profilometry using a Keyence VK-X1000. This method yields two “images” per field of view: a “reflectance” image where pixel intensity corresponds to the intensity of reflected light measured at the detector, and a “height” image corresponding to the height (dimension axial to the objective) of that pixel. Using this, we are able to view both the cell bodies in the reflectance image, as well as their height relative to the underlying hydrogel surface. Cells were manually segmented and tracked and simulation parameters were set to match measurements from these tracks. The experimentally measured parameters include cell length, curvature, width, height, speed, and reversal period.

### B. Simulation

We model *M. xanthus* cells as chains of beads connected by springs with a harmonic restoring force between connected beads as well as on the angles between neighboring springs to model cell stiffness. We use a truncated Lennard-Jones potential to model repulsive interactions between neighboring cells (Fig. 1a). Our simulations are built within LAMMPS which was originally created to model equilibrium systems with precise molecular properties[24]. Additional elements to model out of equilibrium systems with self-propulsive forces and Brownian dynamics were introduced in Winkler et al[25]. These propulsion forces are polar, pointing from bead to bead. We introduced custom functions to orient the propulsion dipoles along the spring connecting each bead to the next bead towards the head of the cell so that the self-propulsion of cells acts along the cell body. We use overdamped Brownian dynamics to model cell motion such that the self-propulsive force drives cells to glide at an intrinsic speed v_0_ in the absence of external forces from other cells, which we measured experimentally. Cells reverse their direction of motion with Poisson statistics and an average reversal period of T which we also measured experimentally. As cells starve and enter their developmental cycle they increase their speed as well as their reversal period. We vary both of these parameters independently and present the experimentally observed changes they exhibit during development as a change in Péclet number, Pe. Péclet number represents the ratio of advection to diffusion and can be calculated according to 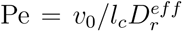, where 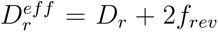 is the effective rotational diffusion constant accounting for cell reversal events[18]. See SI for details of the simulation setup as well as experimental measurements of simulation parameters.

**FIG. 1.**
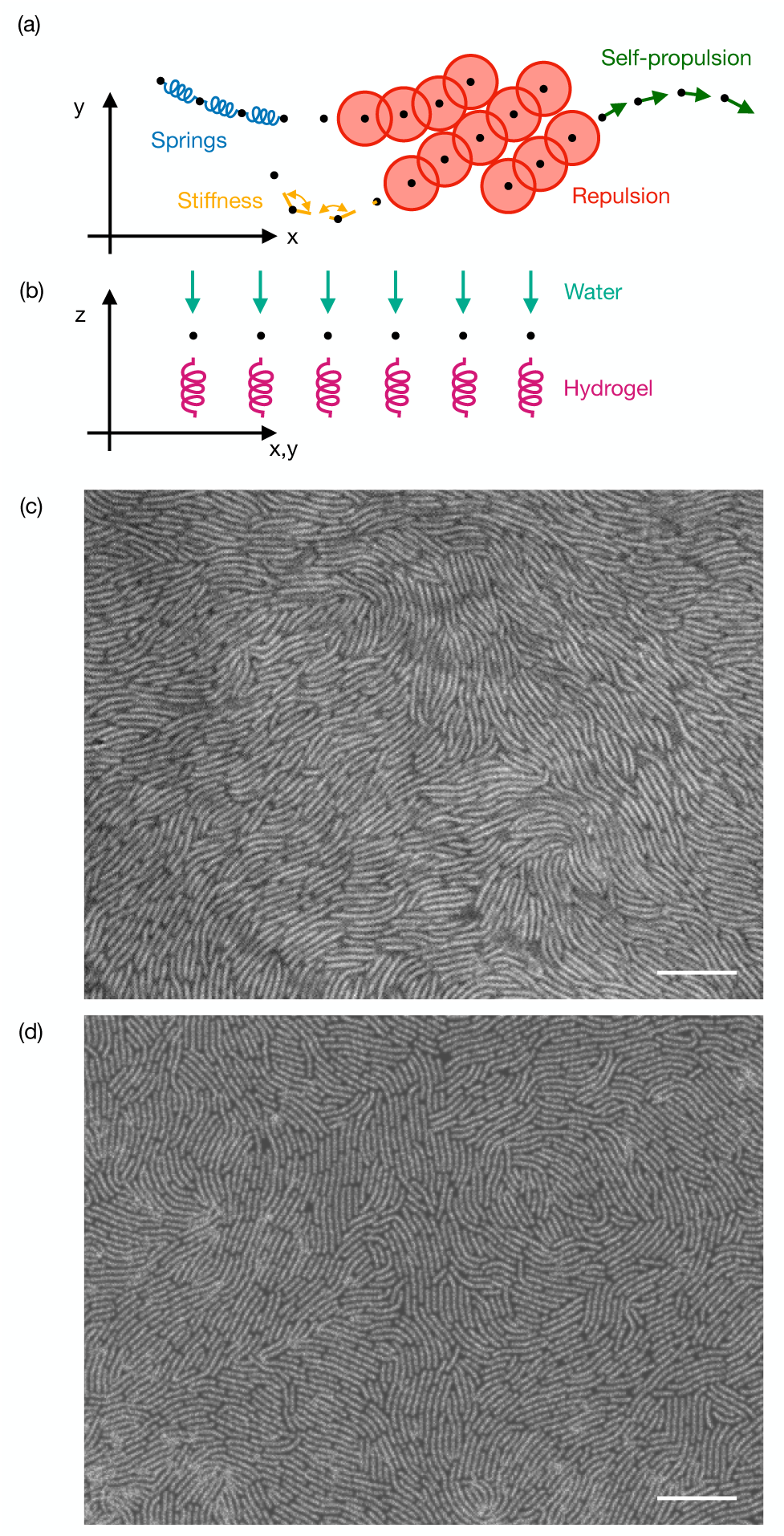
Setup and overview. (a) Simulation schematic, blue springs connecting black bead centers. Red truncated Lennard-Jones repulsive interaction. Yellow cell stiffness, and green self-propulsion. (b) Vertical forces in the simulation, constant downward water force always active shown in teal, and spring upward force towards *z* = 0 which cuts off above that point in magenta modeling the hydro-gel in experiments. (c) Example experiment image, 10*µ*m scale bar. (d) Example simulations frame, 10*µ*m scale bar.

All of the forces in the simulations are fully calculated in 3D. We apply a soft 2D confinement on the cell layer by modeling the hydrogel as an upwards spring that cuts off above z = 0, and the layer of water which coats the cells and holds them on the surface as a constant downwards force (Fig. 1b). This constant downward force does not capture all of the effects of the capillary forces that shape the colony edges and promote nematic order through local pressure[26]. However, in the current paper we aim to study the properties of dense continuous monolayers of cells where these edge effects are minimal, and the computational efficiency of modeling the water layer as a constant downward force allows us to model significantly larger systems. An additional benefit of modeling the water forces as a constant downward force is that we are able to randomly initialize very large numbers of cells in a volume above the surface of the gel and allow them to settle from the constant water forces onto the surface. This gives us the ability to simulate systems at otherwise inaccessible densities with area packing fractions up to or even above one in a satisfying way. This settling process is similar to what occurs in experiments as the inoculation droplet on a hydrogel dries, leaving the cell layer lying flat on the surface. While this initialization method is a useful tool to allow for modeling very high density populations, the details of the initialization do not effect our results as we allow the simulations to run for enough time to reach a steady state before we begin taking data for further analysis.

Most cell parameters are directly measured in experiments such as cell length, width and density. We set the cell bending stiffness in the simulations so that the distribution of cell spline curvatures matches the curvature of cells measured in dense monolayers in the experiments. We finally calibrated the vertical forces in the simulation by measuring a height map of cells on a gel surface using the Keyence profilometer described previously. The height of the top of a cell above the gel surface in simulations is set by the ratio of the gel spring constant and the constant downwards water force. We measure this height experimentally and use it to set the ratio of the vertical forces in the simulations. For the majority of this paper we aim to study the quasi-2D active nematic properties of the cell colony, so we limit cell layering while still allowing for small amounts of vertical motion to relieve local pressure allowing for cells to occasionally cross over each other. To achieve this we set relatively high constraining forces in the vertical direction and scale this force with the cell self-propulsive force. Scaling the vertical forces in this way reduces jamming at low cell speeds allowing them to still relieve local pressure by utilizing the third dimension, while also forcing the high cell speed simulations to remain as a monolayer. The amount of cell layering that we observe experimentally varies widely from experiment to experiment likely due to local variations in drying and levels of excreted polysaccharides. Therefore, scaling the vertical forces in our simulations to maintain a monolayer of cells is not inconsistent with the range of vertical forces also present in experiments.

We run simulations for the equivalent of one hour of experimental time to reach steady state and then analyze simulations results for the following hour, or 6 × 10^6^ time steps. We simulate cells at a density of 0.3µm^−2^ in a periodic box of side length 100µm. As cells are on average 5µm long and 0.7µm wide, this is a cell density with a packing fraction of 1 ± 0.1, as we see in experimental monolayers in high nutrient conditions(Fig. 1c). This simulation box size is similar to the field of view for our microscope which is ∼ 100µm×130µm. Simulated images capture the overall structure of our experimental images, including the formation of a monolayer of densely-packed cells with a high degree of local nematic order and point-like topological defects (Fig. 1d)

## III. RESULTS

### A. Varying cell motility

As *M. xanthus* runs out of nutrients and begins to starve they enter a part of their developmental cycle where cells increase their Péclet number, Pe, (ratio of advection to diffusion) by moving faster and reversing less often [20]. These changes in motility drive a transition from thin layers of cells that spread over a surface to an aggregated state with mounds of cells surrounded by empty space[18]. In our simulations we have independent control of intrinsic cell speed, v_0_, and reversal period, T, without effecting any other cell behaviors. We modulate cell speed by changing the self-propulsion force. Reported cell speeds, referred to here as ‘intrinsic cell speed’ or v_0_, represent the speed at which cells would move in the absence of other external forces, e.g. from surrounding cells in the bulk. We scan over a range for each parameter, covering those displayed by wild type cells naturally during starvation, and measure a variety of properties over this phase space. Fig. 2a shows the changing Péclet number with changing v_0_ and T with the experimentally measured cell speed and reversal period outlined in bold having Pe = 4, and an arrow showing the experimentally observed increasing Péclet number with development. Future plots that are displayed against Péclet number are measured along this development line increasing both cell speed and reversal period simultaneously.

**FIG. 2.**
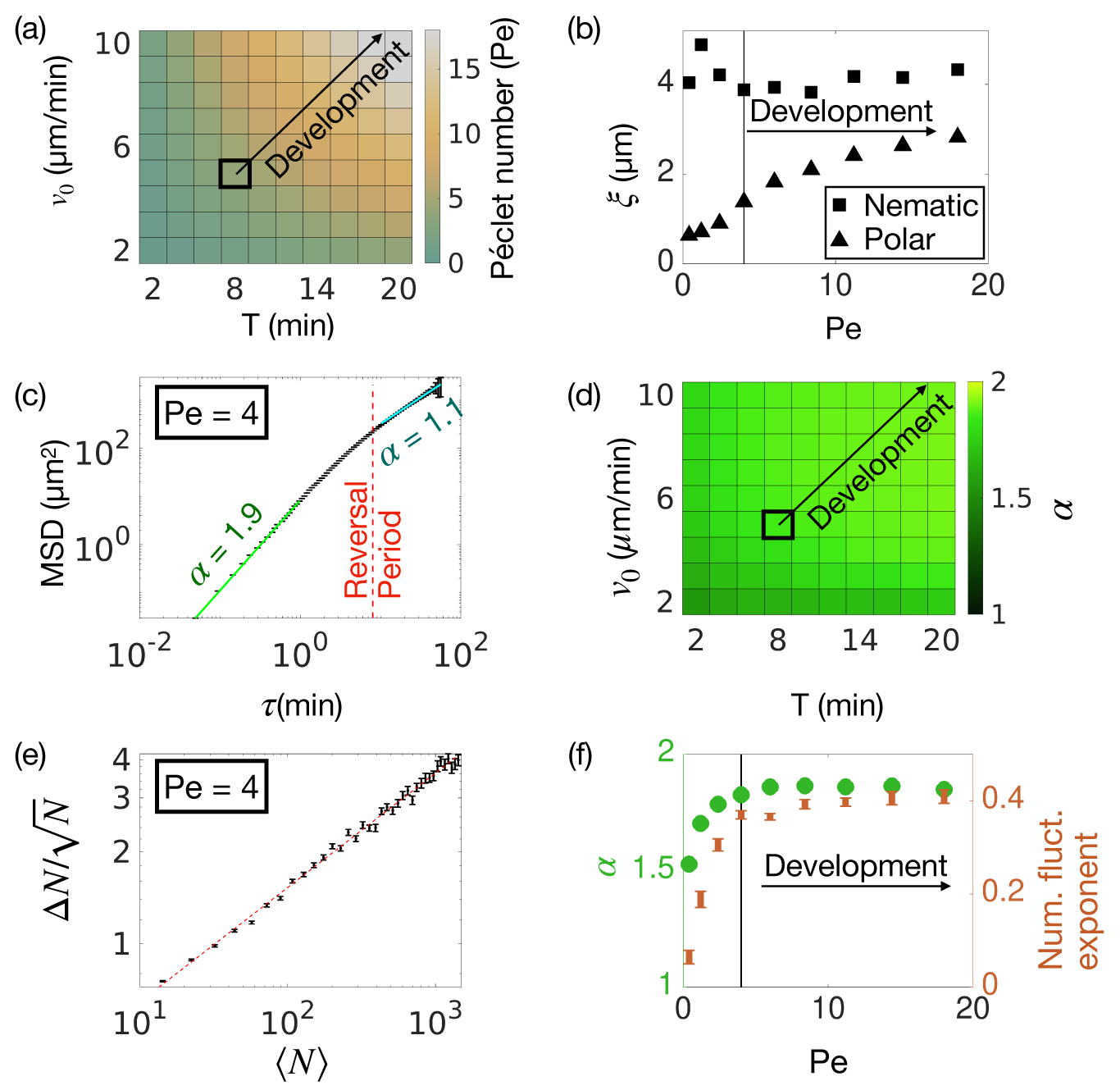
(a) Péclet number, Pe, over a range of intrinsic cell speeds, *v*_0_, and reversal periods, *T* . The value corresponding to experimentally *v*_0_ and *T* is outlined in bold and an arrow shows the direction of motility changes displayed by *M. xanthus* during starvation. (b) Polar and nematic correlation lengths, *ξ* plotted against Pe. The vertical line shows Pe = 4 parameters with the arrow showing the changes made by *M. xanthus* during development. Error bars are the 95% confidence interval for the fit and are smaller than the displayed points. (c) Mean square displacement of individual cells within the dense monolayer at Pe = 4. The green line indicates the fit line for small time scales with *α* = 1.9, while the cyan line shows the fit line for long time scales with *α* = 1.1. The reversal period is indicated by the red line. Error bars are standard error. (d) *α* for the short time scale fits (*τ <* 1min) for cells within a dense monolayer in simulations over a range of *v*_0_ and *T* . (e) Standard deviation of particle number measured for increasing region sizes in simulations at experimentally matched conditions with Pe = 4, plotted against average particle number. Error bars are the standard error. (f) Exponent of number fluctuations calculated from the fit illustrated in (e) plotted against Pe during development (orange) along with the short time scale *α* values extracted from the phase space in (d)(green). Error bars are the 95% confidence interval for the fit lines, and are smaller than the plotted points for *α* values.

We measure the organization of cells within dense monolayers by measuring both the nematic and polar correlation lengths, ξ, (Fig. 2b). The polar correlation length reflects the domain sizes over which cells align their self-propulsion direction, while the nematic correlation length reflects the domain sizes over which cells align their long axes with 180^◦^ symmetry independent of their head/tail orientation. We find that the nematic correlation length is always greater than the polar correlation length indicating that the nematic domains are larger than polar domains suggesting that these cell monolayers form a nematic material. Interestingly the polar correlation length increases during development while the nematic one remains constant.

To examine how cells move within the established nematic structure of these dense monolayers we measured the mean square displacement, MSD, of cells within the system at Pe = 4 (Fig. 2c). Plotting the MSD on a log-log plot and fitting a line we can extract the diffusivity exponent, α, as the slope of the fitted line in log-log space. We find that at short timescales α is close to two, suggesting that cells move ballistically and are able to glide past each other uninhibited. At timescales above the reversal period this changes to a value of α close to 1, which represents purely diffusive motion, and is the same trend observed previously in low density reversing self propelled rods[8]. We measure α at small timescales (τ < 1min) over our phase space to examine where cell motion is uninhibited (Fig. 2d) and find that cells move ballistically at short timescales (α ∼ 2) for the majority of the phase space. However, for low v_0_, the small timescale αs drop significantly, indicating that cell motion is inhibited in this regime. This inhibition is likely due to the decreased average active force in this regime causing cells to be unable to overcome the vertical confining forces and release local pressure, something reminiscent of a jamming transition (see SI2).

As an active material we expect our system to display giant number fluctuations[27]. We measure this by breaking the system into smaller boxes of different sizes, and measure the average (⟨N⟩) and standard deviation (ΔN) of the number of beads in the all boxes of increasing sizes. Normalizing ΔN by 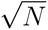 and plotting it on a log-log scale against ⟨N⟩ will result in an exponent of 0.5 for an active material and 0 for a passive one, and indeed we observe giant number fluctuations with an exponent of 0.5 as expected for an active material at our experimentally measured parameters(Fig. 2e)[27]. During development, with increasing Péclet number, cell motion remains uninhibited and the colony forms an active material as seen by α remaining close to 2 and the number fluctuations exponent close to 0.5. However, interestingly, for very small Péclet numbers the cell motion is inhibited and the system forms something more like a passive nematic liquid crystal(Fig. 2 f). Having found that our cells layers form an active nematic material, we set out to measure defect properties within the cell monolayers during development.

### B. Topological defects

Topological defects are locations within a nematic material where the molecules become misaligned, and they have been shown to be an important feature of active nematic materials. In the absence of topological defects particles are well aligned and local active stresses cancel out. However, in an extensile active nematic material, local bend deformations in particle alignment are unstable and grow leading to the formation of pairs of oppositely charged defects[28]. These topological defects provide sites where the local active stress drives flows of material leading to comet shaped +1/2 defects self propelling while the threefold symmetric −1/2 defects diffuse passively[29]. The self-propelling +1/2 defects can provide a mechanism for mixing within active nematic materials[30] and the flows of many defects combine to cause the material flows to form vortices and exhibit turbulence[31].

We identified topological defects by interpolating the bead dipoles, which point along the cell bodies, to find the orientation of the director field (θ). We then calculate the winding number 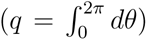 on a grid at every vertex of four interpolated grid points. We then combine all charged points within a minimum distance of each other, r*_min_* = 1.4µm (two cell widths), by summing their charges and averaging their locations in order to filter out transient local charge pairs that arise due to small fluctuations in bead orientations and noise. This also allows us to detect any integer charge defects if they should arise, though we very rarely if ever detect them in our simulations or experiments. We observe ±1/2 defects in our simulations which closely resemble those seen in experiments. The examples in Fig. 3a (experiments) and Fig. 3b (simulations) show a cell image overlayed with local cell orientation, or director field (θ), along with a defects pair with the +1/2 defect displayed as a red comet shape, and −1/2 defect as a blue triangle.

**FIG. 3.**
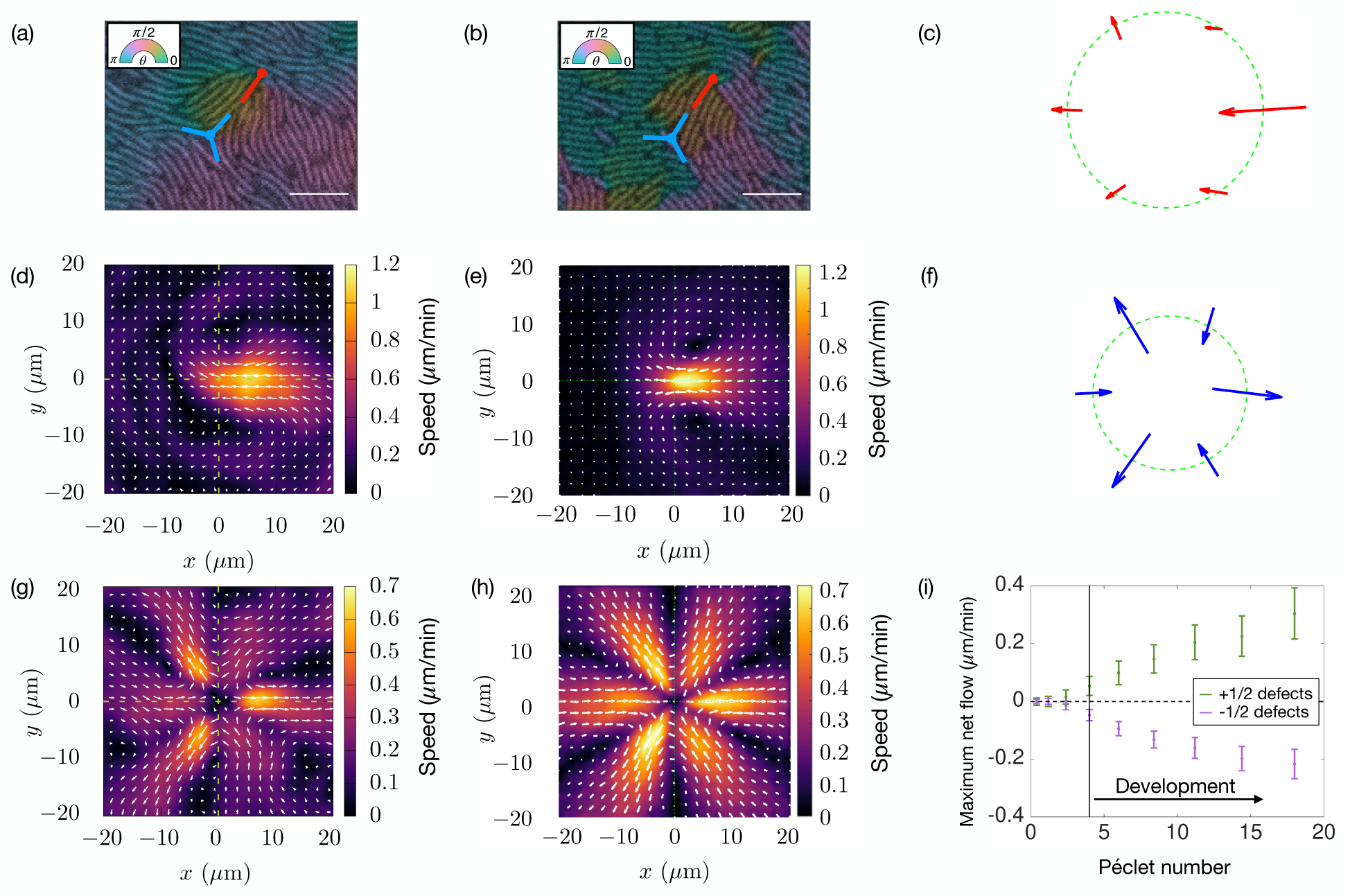
Defects and flows. (a) A pair of defects from an experimental image overlayed with color representing the local director, 5*µ*m scale bar. (b) A pair of defects from simulations also colored by local director, 5*µ*m scale bar. Positive defects illustrated in red, and negative in blue. (d, g) Experimental flow fields averaged over all positive and negative defects respectively. (e, h) Simulated flow fields averaged over all positive and negative defects respectively. (c, f) Illustration of the velocity vectors at the radius where the magnitude of the net flow is maximized for positive and negative defects respectively, the net flow is calculated as the sum of the radial components of the displayed vectors. (i) The maximum net flow at positive and negative defects measured against Pe.

Next we measured the average flow field around each type of defect. In the experiments we use the Farneback algorithm to measure optical flow across the whole field of view at all times, while the simulations simply output the velocity vector for each bead at each time point. From these individual velocity flow fields we extract and reorient the local velocity field around each defect so that all defects are centered and aligned. We then average together the flow fields for each charge of defect to obtain a mean velocity field around +1/2 defects in experiments (Fig. 3 c), and simulations (Fig. 3 d), as well as around −1/2 defects in both experiments (Fig. 3 f) and simulations (Fig. 3g). The flow fields around defects in simulations share the same structure and are very similar even in magnitude to those measured in experiments despite the fact that the individual cell speeds are nearly five times higher (∼ 5µm/min) than the peak speed in the average flow fields(∼ 1µm/min), confirming that the simulations are capturing the basic mechanisms and mean-field defect behaviors observed in experiments quantitatively as well as qualitatively.

Cell aggregation which leads to fruiting body formation is driven at this intermediate length scale by the accumulation of cells at +1/2 defects and depletion from −1/2 defects [22]. We measure the amount of accumulation or depletion taking place by measuring the net flow through a circle around each type of defect at a radius (R) which maximizes the magnitude 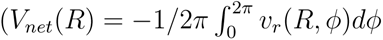, where 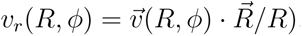. Fig. 3 (c,and f) shows a schematic taken from the simulation flow fields, illustrating the flow through a circle around the center of +1/2 and −1/2 defects respectively, the net flow is the sum of the radial component of these flows, with the net flow plotted in Fig. 3 taken at the radius which maximizes the magnitude of this quantity. Finally we calculate the net flow around positive and negative defects against Péclet number and find that the magnitude of the net flow around both charges of defects steadily increases with development. The negative values for −1/2 defects reflects a net outflow while the positive values represent a net inflow seen at +1/2 defects. Knowing that the average inflow or outflow increases around defects with development we next measure what the defects themselves are doing.

### C. Defect dynamics

As cell accumulation and depletion occurs at topological defects, not only will the net flow effect this accumulation process, but defect density and motility will also likely play a role. We expect defects which remain stationary for longer to allow for greater accumulation at single locations over time, compared to defects which are too motile[5]. To measure defect motility, we track defects over time by simple nearest neighbor tracking between frames while conserving charge, and use these defect tracks to calculate the mean square displacement over time with Pe = 4 (4a). As expected for an active nematic material, where +1/2 defects self propel, the slope of the MSD on a log-log plot for +1/2 defects is greater than that of the −1/2 defects which are expected to be more diffusive[29]. We then fit the MSD for +1/2 and −1/2 defects and measure α vs. Pe during development and find that α is always higher for the +1/2 defects than −1/2 ones. However, they both remain relatively constant or perhaps very slightly increasing(Fig. 4b). This suggests that during development defect motility is mostly constant. However, at very low Pe number where the bacteria colony appears more like a passive nematic material than an active one, α drops to something close to 1 and is roughly equal for both positive and negative defects as expected in a passive nematic liquid crystal.

**FIG. 4.**
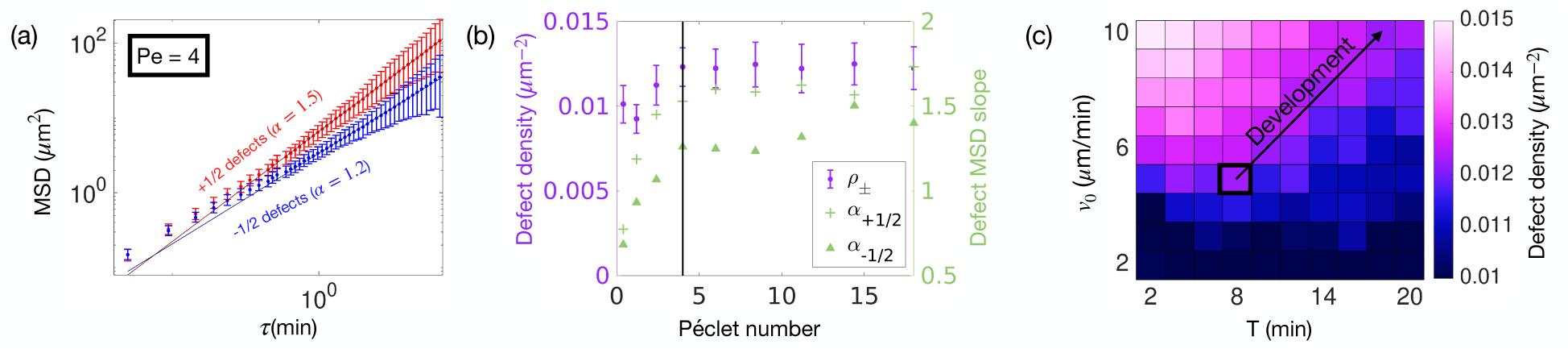
Defect behaviors. (a) Mean square displacement of positive (red) and negative (blue) defects for experiment matched conditions. (b) (purple, left axis) Defect density against Pe with standard error as the errorbars. (green, right axis) Fit exponent (*α*) for positive (plus signs) and negative (triangles) defects plotted against Pe during development. The error bars as the 95% confidence interval from the fit are smaller than the width of the line. (c) Phase space of defect density against both *v*_0_, and T. Experimentally matched conditions outlined in bold and an arrow representing changes displayed by cells during development.

We then measured the average defect density across all frames and simulations to examine how it changes with development. We find that defect density remains constant with increasing Pe during development(Fig. 4b). Interestingly, if we examine defect density across the entire phase space of v_0_ vs. T we find that the relationship of defect density with Pe is not as simple as it might first appear (Fig. 4c). The fact that cells increase their Péclet number by changing both speed and reversal period simultaneously is necessary for the defect density to remain constant. If cells were to only increase their reversal period without also increasing cell speed the defect density would decrease. Conversely, if cells increased speed without increasing reversal period the defect density would actually increase.

Together these results reveal that as cells increase their Péclet number by both increasing their speed and reversal period during development, they maintain a constant nematic and defect structure within the cell colony.

### D. Cell aggregation

As cells speed up and reverse less often the colony aggregates and begins to pile up, causing it to have regions with two layers of cells as well as areas with no cells at all. This presents challenges for accurately simulating the cell layers as the details of the interactions between the first and second layer of cells is unclear. Cell gliding is driven by motors which form focal adhesions with the surface they are gliding across and how these gliding forces are transmitted when cells are gliding on top of other cells is unclear[32]. To avoid these complications in the previous sections of this paper we scaled the vertical confinement forces with the active force in order to prevent layering. In order to see whether the aggregation in our simulations is consistent with that observed experimentally, we now fix the vertical confining forces to an intermediate value while varying only cell speed and reversal period. Fig. 5(a) shows an example simulation output image at our experimentally measured condition in high nutrients, with Pe = 4, where we expect cells to spread out to cover the surface, and indeed we see that they do. Fig. 5(b) shows an example image at Pe = 18 with higher speed and reversal rate as seen in experimental systems when cells begin to aggregate into fruiting bodies during starvation. We see a similar aggregation behavior in our simulations with both an increase in empty areas as well as regions with two layers of cells.

**FIG. 5.**
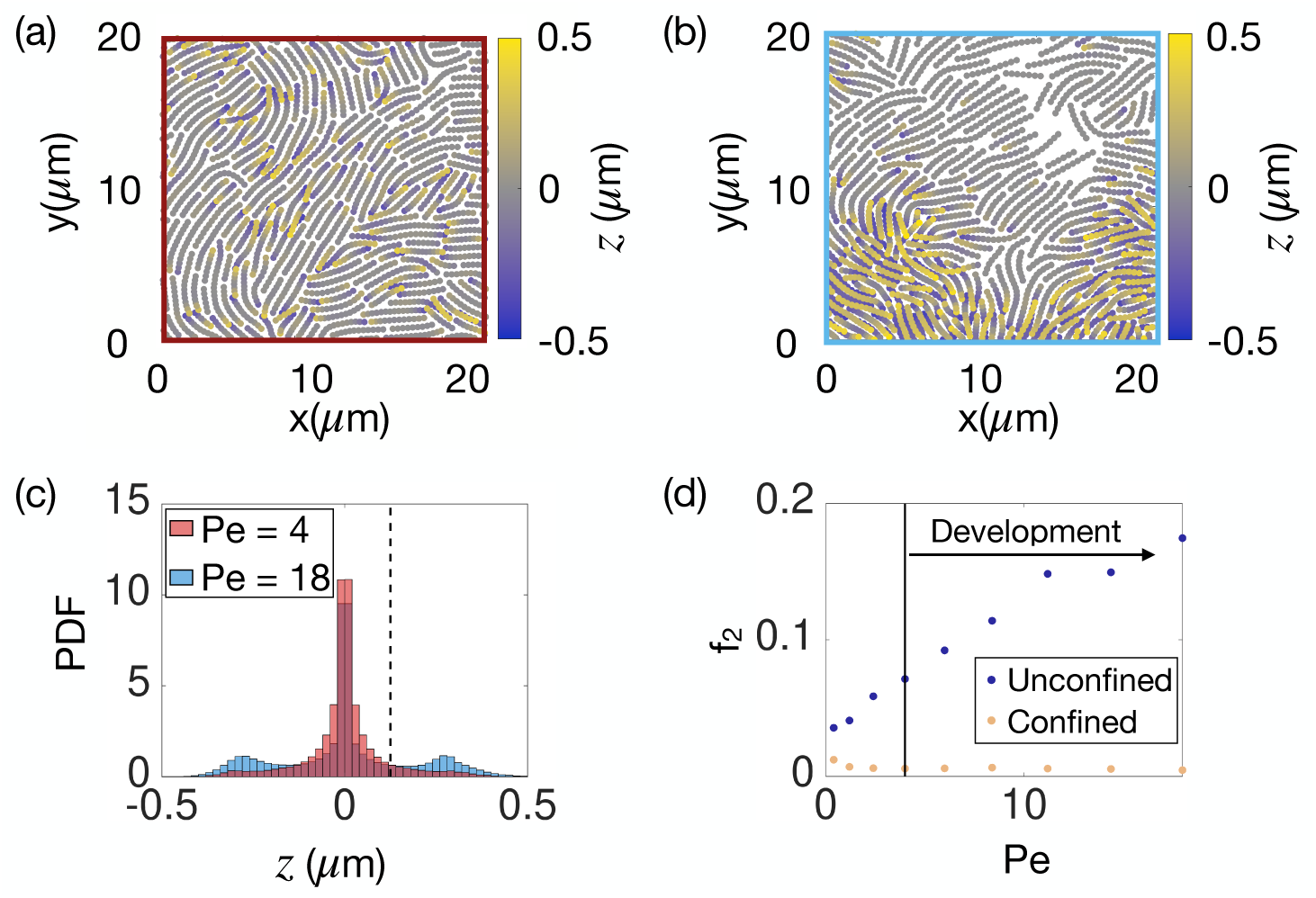
Aggregation. (a, b) Example simulation output colored by bead *z*-position. (a) Parameters are set to the experimentally measured high nutrient conditions with Pe = 4 where a monolayer is expected. (b) Parameters are set to high speed and reversal period with Pe = 18 where aggregation and increased layering is expected. (c) Histogram of bead *z* positions for the two examples shown in (a, Pe = 4) and (b, Pe = 18). The dashed vertical line shows the height threshold between the bottom and top layer of cells. (d) The fraction of cells in the second layer plotted against Pe during development. The peach points show the simulations where the vertical z forces are scaled with active force in order to prevent layering and maintain a monolayer of cells as presented in the previous sections of this paper. The blue points show the values when the vertical forces are held constant at an intermediate value while only cell speed and reversal period are varied. Error bars are the standard error and are smaller than the displayed points.

We quantify the amount of aggregation and layering occurring by simply looking that the z positions of the simulated cells. Fig. 5(c) shows a histogram of the z position of beads for the two images shown in (a) and (b). We can see that for the low Pe conditions (red) there is a single central peak in the histogram at a value of zero illustrating that the cells form a single monolayer in these conditions. However, at high Pe we see this same central peak as well as two side peaks. The central peak is from the regions of the simulations where a monolayer exists while the two side peaks are from regions with two layers of cells. Since the hydro-gel is modeled as a simple harmonic spring force, when cells climb into the second layer they also push the first layer of cells down into the gel resulting in a value of z ∼ 0.3 for cells in the second layer, and z ∼ −0.3 for cells in the layer below them. These height values are consistent with cells of a radius of 0.7µm interweaving within the grooves between adjacent cells in the bottom layer. This process of cells climbing up into a second layer as well as being pushed down into the gel are both consistent with previous experimental observations where second layers of cells on stiff gel substrate emerge above the bottom monolayer of cells [22], while cells aggregating on a very soft gel actually form the second layer downwards within the gel as this presents less resistance than deforming the water layer that coats the colony[5, 26].

To examine this aggregation trend with increasing Pe during development we measure the fraction of beads with z > 0.15µm, which are those in the second layer, and plot this fraction against Pe. Fig. 5(d) shows this fraction during development when cells are free to pile up (blue), and indeed we see an increase in fraction of cells in the second layer with increasing Pe, indicating an increase in aggregation during the development process as expected. To confirm these calculations we also measured the fraction of cells in the second layer for the simulations from the previous sections where cells were prevented from climbing into the second layer in Fig. 5(d, peach). Indeed, in these conditions we see that cells are unable to pile up or aggregate as we designed it to be.

## DISCUSSION

*Myxococcus xanthus* is a soil bacteria with a developmental cycle where large collectives of hundreds of thousands to millions of cells transition from a thin spread out sheet of cells and aggregate into 3D droplets of cells separated by empty space. We know that this aggregation process is driven by individual cells tuning their motility to increase Péclet number (Pe) by speeding up and reversing less often[18]. We also know that on an intermediate length scale cells tend to align their long axes within a local neighborhood and the 2D to 3D transition is initiated by layering events at topological defects where cells are misaligned[22]. Here, we present a simulation of *Myxococcus xanthus* as bead-spring chains where we can investigate the properties of topological defects as cells increase their Pe.

We found that modeling *M. xanthus* populations as a dense collection of flexible self-propelled rods, or bead-spring chains, captures the macroscopic monolayer structures observed in experimental systems with the same topological defects present. Varying the Pe in simulations by increasing both cell speed and reversal period drives the system into an aggregated state just as the cells do naturally in experiments. Looking closer at the cell structure within the system with increasing Pe we found that firstly the nematic correlation length is always longer than the polar one suggesting that we have a nematic material. While the nematic correlation length remains constant with Pe, the polar correlation length actually increases with increasing Pe suggesting that any effects driven by instantaneous cell polarity may increase during development.

Within the nematic structure of the cell monolayers, cells are able to glide past each other uninhibited by their surroundings, as indicated by the short timescale αs of cell mean square displacements remaining close to 2. The number fluctuations scale with mean particle number with an exponent greater than 0 confirming that we have an active nematic material. However at very small Pe both α and the number fluctuations exponent drop significantly suggesting that when cell motion is inhibited by surrounding cells in a dense monolayer, the material begins to behave more like a passive nematic than an active one.

When we measure the flow around topological defects during development we find that there is an increased inflow at positive defects with increasing Pe, and a corresponding increased outflow at negative defects, which drives aggregation during starvation. Investigating the motion of defects and their density reveals that these quantities do not change significantly during development. Interestingly, the constant defect density relies on cells modulating their Pe by changing both speed and reversal period simultaneously. Were cells to only increase speed or reversal period during starvation and not both, the defect density would also change with starvation rather than remaining constant.

In this paper we showed that a dense bead-spring chain model captures the collective behaviors of *M xanthus* bacteria colonies. As cells starve and increase their Pe they maintain an unchanging nematic structure while increasing the net flow around topological defects driving the aggregation process. This increased net flow may be tied to an increasing polarity we also observe with increasing Pe. The question of why a constant defect density may be advantageous during starvation is unclear and a topic for future study, as is the long term aggregation process connecting this initial layering to the full formation of relatively large 3D droplets.

## AUTHOR CONTRIBUTIONS

KSC designed and ran simulations. KC and MB performed and analyzed experiments. JWS supervised the study. All authors interpreted data and results. KC wrote the manuscript with input from all authors.

## DATA AVAILABILITY

Simulation code including custom functions added to lammps is available at: https://github.com/kcopenhagen/lammps-myxosim.

## ACKNOWLEDGEMENTS

We thank N Wingreen, C Fei, A Bourque, B Figueiredo, A Al Harraq, R Alert, and A Pyo for discussions. This work was supported in part by the National Science Foundation, through award PHY-1806501 and the Center for the Physics of Biological Function.

## Appendix A Model parameters

We measure model parameters from experimental data of dense monolayers of *M. xanthus* as described in the methods section. We manually segmented and tracked cells to measure parameters which are then rounded for simplicity when plugged into the simulations with the values displayed in Table I. Distributions and details of parameter measurements are presented below.

**TABLE I.** Table of simulation parameters including value and a brief description of the parameter.

| Parameter | Value | Description |
| --- | --- | --- |
| w | $0.7\mu\text{m}$ | Cell width. |
| $K_{stiff}$ | $80\text{kg}\mu\text{m}/^\circ\text{min}^2$ | Cell stiffness. |
| L | $5 \pm 2.5\mu\text{m}$ | Cell length. |
| $\rho$ | $0.3\mu\text{m}^{-2}$ | Cell density. |
| $v_0$ | $2 - 10\mu\text{m}/\text{min}$ | Intrinsic cell speed. |
| $T_{rev}$ | $2 - 20\text{min}$ | Reversal Period. |
| $F_{H_2O}/K_{gel}$ | $0.3\mu\text{m}$ | Ratio of vertical confinement forces. |
| $K_{gel}$ | $100 \times v_0 \text{ kg}/\text{min}^2$ or $500\text{kg}/\text{min}^2$ | Spring constant of the gel. |
| $K_{spr}$ | $500\text{kg}/\text{min}^2$ | Spring constant of springs connecting beads. |
| $r_{spr}$ | $0.4\mu\text{m}$ | Spring length. |
| $\epsilon$ | $5\text{kg}\mu\text{m}^2/\text{min}^2$ | Lennard-Jones well depth. |
| dt | $2 \times 10^{-5}\text{min}$ | Simulation timestep. |

### 1. Cell Shape

Cell shapes were measured from analysis of the bright field images as seen in Fig. 1(c).

Eight experiments were analyzed as described here.

#### a. Cell Width

We measure the cell width, or more accurately the spacing between neighboring cells, by taking a 2D fourier transform of the experimental laser brightness image. Binning and averaging by radius reveals a sharp peak at the cell spacing value (Fig. S1(a)). We measure the distribution of this cell spacing across all frames in all experiments and use this spacing to set the bead diameter in simulations (Fig. S1(b)).

#### b. Cell length and stiffness

We obtain cell masks for the cells in experimental dense monolayers by manually tracing cells from experimental images of monolayers (Fig. S2a,b). We manually traced cells from 19 images with similar appearance to those shown here. We then skeletonize the cell masks and generate a cell spline along the cell body center lines (Fig. S2c). Using these cell splines we measure a distribution of cell curvatures by measuring a distribution of the angles between sets of 3 connected spline points as seen in black in Fig S2d. We ran simulations with a range of cell stiffness constants and calculated this same angle distribution by converting simulation outputs into cell masks and using the same protocol as we did for experimental data to extract splines and calculate angle distributions. We selected the cell stiffness as the value which resulted in simulated cells that most closely match the curvature of the experimental cells. We initialize each cell as a string of beads connected by springs in a chain where the number of beads determines cell length which is selected from 5 ± 1µm with a gaussian distribution based on the experimentally measured cell lengths shown in Fig. S2e.

We set the simulation cell density to have an area packing fraction of 1. For our cells with average cell length of 5µm and width of 0.7µm, this results in an approximate cell area of A = 5 × 0.7µm^2^. To have a packing fraction of 1 we set the density to ρ = 1/A = 0.3µm^−2^. The images for manual tracing were selected to be only those without any double layers in order to be able to distinguish individual cells. However, we measured the area density of cells in our manually traced cell masks anyways and found values close to but slightly lower than the 0.3µm^−2^ used in simulations. This is expected given the biased images selected for cell tracing with slightly lower density. Therefore, the density value used for simulations is consistent with the numbers measured experimentally.

### 2. Cell Motility

We measure cell motility parameters by manually tracking cells from frame to frame in experimental monolayers. We tracked at least 10 cells from each of the 8 experimental data sets with a total of 120 cell tracks and extracted cell speeds and reversal periods from these tracks. The experimentally measured values are shown below however both of these parameter are varied in the simulations as they are properties that cells tune internally during starvation.

#### a. Self Propulsion

From our manual cell tracks we calculate instantaneous cell speeds between sequential frames. The self-propulsion force in simulations drives cell gliding so we use the speed distribution measured experimentally to set the propulsion force shown in Fig. S3 (a).

#### b. Reversal Period

We identify reversal events in cell tracks manually as well and calculate a distribution of times between reversal events to set the reversal period in simulations S3 (b). The experimentally measured reversal period is fit with an exponential distribution with a delay offset that is fit as well due to minimum detection limit between reversal events. In simulations we simply use a Poisson process for reversals with the average determined by this fit.

### 3. Vertical forces

We estimate the vertical forces in simulations by using the height measurements of cell layers in experiments in the form of a topological height map. The vertical forces consist of an upwards spring force which cuts off above z = 0 modeling the agar gel surface, and a constant downward force modeling the water layer which coats the cell colony holding it onto the surface. The equilibrium height of cells is determined by the height at which these forces are equal, −K*_gel_* × Δz = F*_H_*_2_*_O_* . Therefore, we can determine this ratio by measuring the distance between the gel surface and the top of the cells in experiments. Fig. S4 (a) shows the distribution of heights for each experiment. The small peaks at zero are from regions where there are holes in the monolayer and no cells are present. The majority of the experimental images are areas with a single monolayer of cells so the the large peak in the height distribution shows the height that the top of the cells sits above the gel surface. This peak is the height that corresponds to the ratio of vertical forces (F*_H_*_2_*_O_* /K*_gel_*). The location of the peak varies from experiment to experiment likely due to slight differences in local gel stiffness and water content due to drying which was not controlled in these experiments. We used an intermediate value of 0.3µm as this ratio when setting the simulation parameters.

The magnitude of these forces sets how easy it is for cells to pile up into second layers. With the ratio of F*_H_*_2_*_O_* /K*_gel_* fixed we varied the magnitude of both forces to investigate reasonable values to use in simulations. We measured the area fraction of cells that are a single monolayer and compared this to an estimate of this quantity measured from simulations over a range of vertical confinement forces. The experimental measurements cover a wide range across all experiments and we are able to vary K*_gel_* in simulations to cover the whole range seen experimentally as shown in Fig S4 (b). In the first half of this paper we let K*_gel_* scale with the cell self propulsion which both prevents cells piling up at high cell speeds and jamming at low cell speeds allowing us to study quasi-2D properties of the cell monolayers. In the last part of this paper we fix K*_gel_* = 500kg/min^2^ to allow cells to aggregate and pile up for comparison with cells during development when aggregation occurs experimentally.

### 4 Additional Parameters

#### a. Bead springs

The springs connecting beads simply act to hold the cell lengths so we set this parameter (K*_spr_*) sufficiently large in order to overcome all other forces in the simulations and prevent cells becoming compressed or stretched. The spring length or bead spacing was set to 0.6×w so that there is some overlap between neighboring beads within a single cell resulting in a somewhat smooth cell outer surface.

#### b. Repulsion

The Lennard-Jones well depth sets how strong the repulsive forces are between neighboring cells. When repulsive forces are too high the beads of neighboring cells can become interlocked and cells are unable to move freely at high densities. We selected a Lennard-Jones well depth sufficiently below this threshold while still being high enough to prevent cells passing through each other. There is a large range of values that satisfies these conditions and provided the well depth falls within this range the exact value does not effect our results.

#### c. Time step

Provided the simulation time step is small enough to prevent numerical integration errors, our results are independent of time step. Smaller time steps will in theory give better results but increase simulation run time so we selected a somewhat arbitrary time step and simply ensured that it falls below a value that could cause any numerical integration errors or losing beads.

## Appendix B Physical cell inhibition

In main text Fig. 2(f), we observed that the α values extracted from single cell MSDs were approximated equal to 2 for high Pe indicating uninhibited self-propulsion or ballistic motion. However, at low Pe, α decreased to something between one and two where a value of one would indicate purely diffusive motion. This decrease could be either due to a shorter timescale turn over from ballistic to diffusive motion in the MSD, resulting in a lower slope from the fit, or due to cell propulsion becoming inhibited or blocked in this regime. We believe the cells are becoming inhibited for several reasons. The first is that the fits were performed on the MSD for time windows below 1min which is significantly lower than even the shortest reversal periods so the ballistic to diffusive transition should not effect these fits. We also examined if this decreased α value could be due to inhibition or being crowded and blocked by other cells in the dense layers with something reminiscent to a jamming transition. To test this we compared the single cell MSDs and extracted α values from the simulations with fixed z confinement forces across the phase space Fig. S5(a). In these conditions the vertical confinement forces when v_0_ < 5 are larger than for the phase space shown in main text Fig. 2(d). With these larger confining z forces we find that the α values drop significantly more for low v_0_ when when compared to Fig. 2(d). As cells glide on the surface they occasionally relieve internal pressure in the cell layers by temporarily pushing into the third dimension. This can be seen by some cell ends pushing up or down slightly in Fig. 5(a). Cells are only able to relieve local pressure with this method if the self-propulsion force can sufficiently overcome the vertical confining forces. Therefore, when the confining forces are higher at low cell speeds the cells are unable to relieve local pressure and we see a transition reminiscent of jamming where α drops and cell motion becomes inhibited through crowding and trapping by neighboring cells in this regime.

**Fig. S1.**
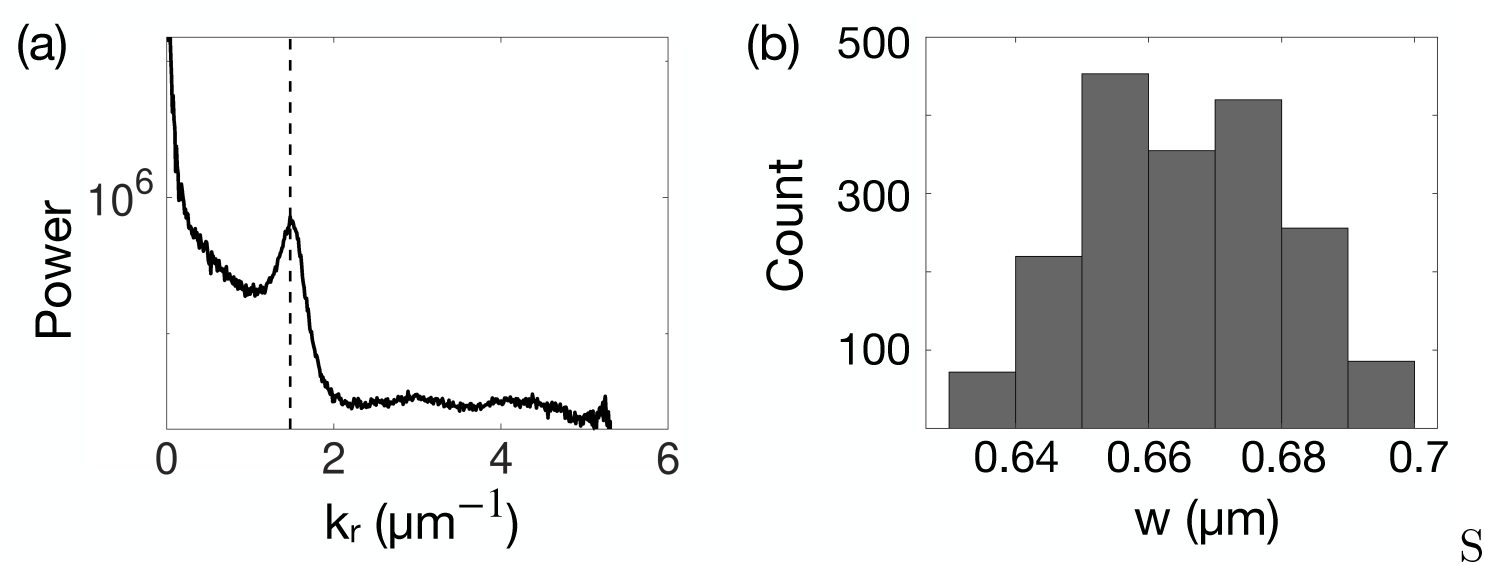
(a) Radial power spectrum from a single representative laser brightness image from experiments. The peak in the power spectrum is highlighted by the vertical dashed line. (b) Distribution of cell widths extracted from the peaks of power spectra from all frames in all experiments.

**Fig. S2.**
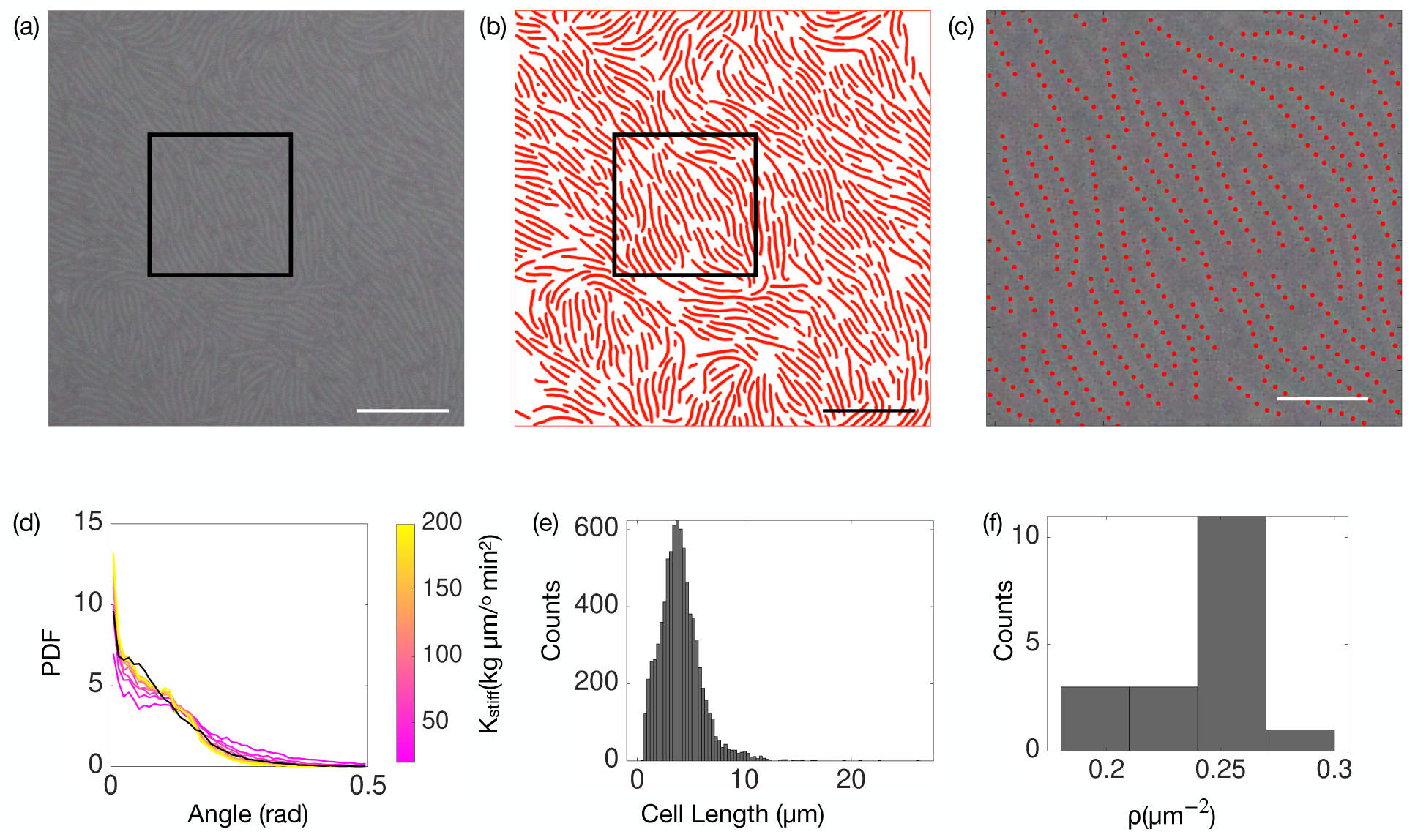
(a) Experimental bright field image of a monolayer of cells. Scale bar is 10*µ*m. (b) Manually traced cells from the image in (a). Scale bar is 10*µ*m. (c) A zoomed in view of cell splines extracted from cell masks within the box drawn in (a) and (b) overlayed on the experimental brightness image. Scale bar is 3*µ*m.(d) The black line is the distribution of angles between sets of 3 consecutive spline points from all manually traced experimental cells. The colored lines are the distribution of sets of 3 beads from simulations colored by a range of stiffness parameters displayed in the colorbar. (e) Distribution of cell lengths. (f) Distribution of density in manually traced cell monolayers.

**Fig. S3.**
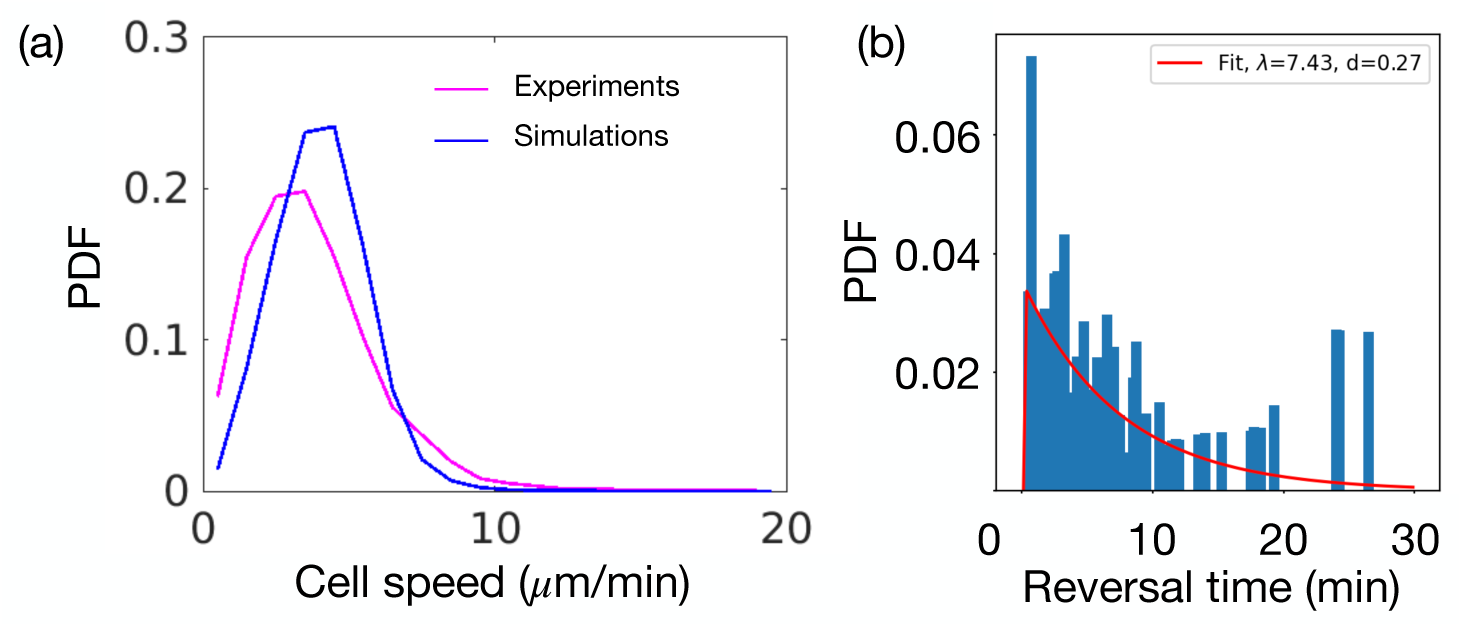
(a) Distribution of cell speeds from manually tracked cells in experiments and the distribution of values used in simulations. (b) Distribution of reversal periods extracted from manually traced cells with an exponential fit with a delay period which is fit as well.

**Fig. S4.**
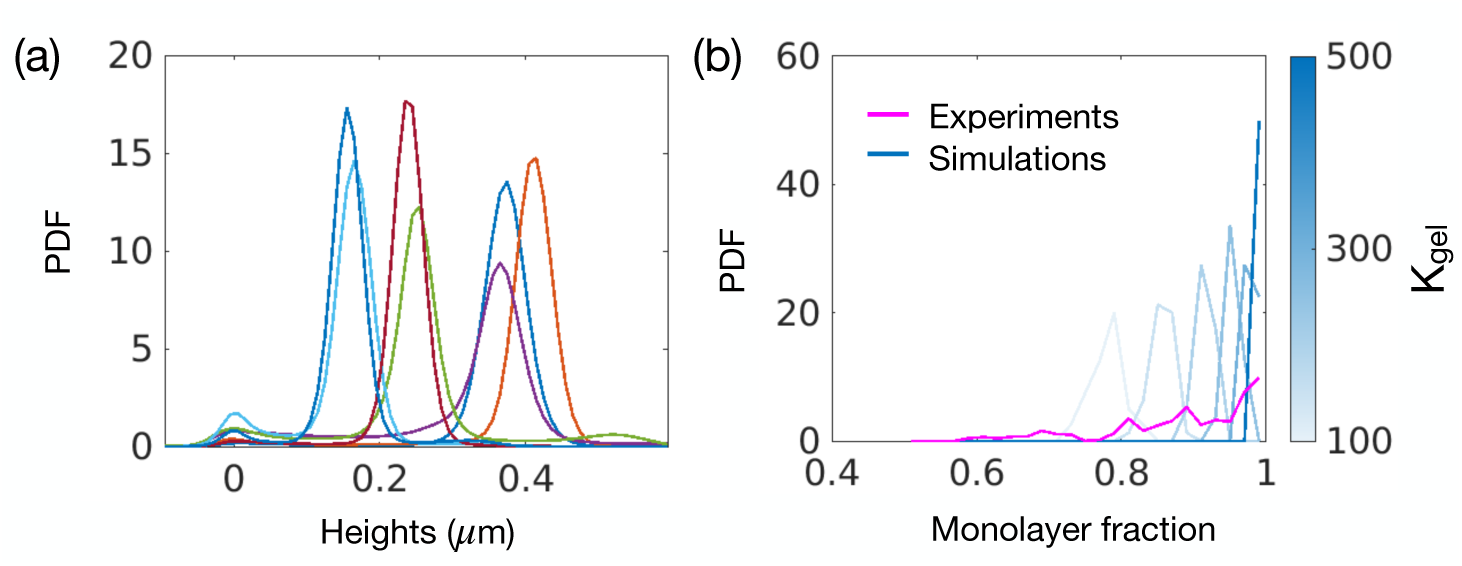
(a) Distribution of experimentally measured heights across the surface of the cell layers. Each color represents a separate experiment. (b) Distribution of the fraction of the area within the field of view that are a single monolayer (no holes or second layers). Experimental distribution shown in pink and simulations in blue for a range of gel spring constants shown by the color.

**Fig. S5.**
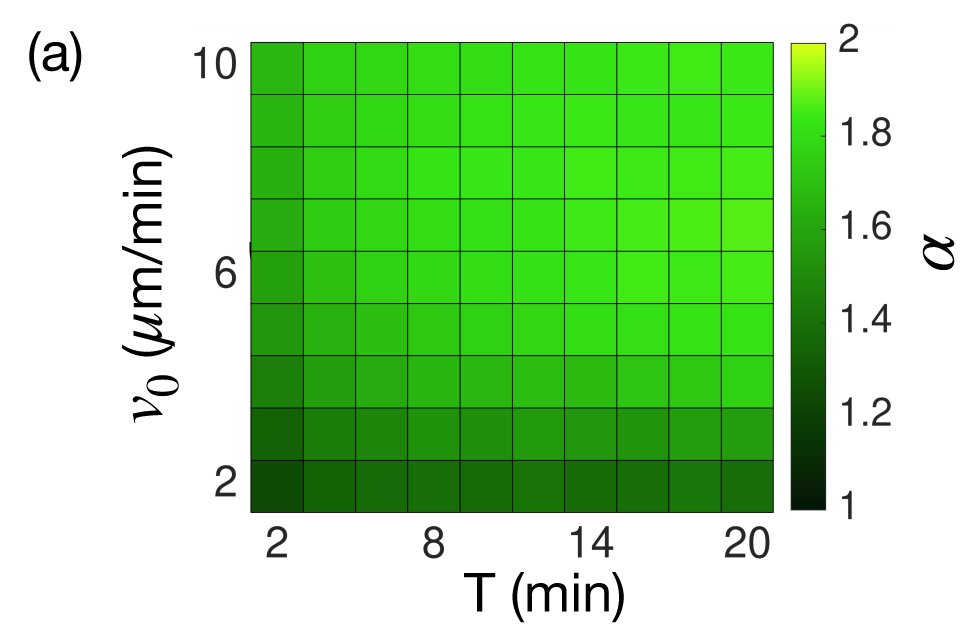
(a) *α* extracted from single cell MSDs for the simulations where the vertical forces are fixed across the entire phase space.

